# Disease progression is associated with differential neutrophil maturation in *Mycobacterium tuberculosis-infected* macaques

**DOI:** 10.1101/2025.06.23.661022

**Authors:** Stanley B. Dinko, Candie Joly, Paul Mazet, Gaëlle Sandillon, Victor Magneron, Natalia Nunez, Céline Mayet, Ségolène Diry, Cassandra Gaspar, Marco Leonec, Sophie Luccantoni, Camille Ludot, Benoit Delache, Emma Jougla, Julie Morin, Wafa Zouaoui-Frigui, Roland Brosch, Vanessa Contreras, Anne-Sophie Galloüet, Nathalie Bosquet, Francis Relouzat, Quentin Pascal, Baptiste Jean, Marion Holzapfel, Olivier Lambotte, Thibaut Naninck, Roger Le Grand, Julien Lemaitre

## Abstract

Tuberculosis (TB), caused by Mycobacterium tuberculosis (Mtb), is associated with clinical diversity and outcomes ranging from latent TB to active TB with distinct pathophysiologies. However, our understanding of the innate immune mechanisms related to the protection or progression of TB is limited. Among innate immune cells, the role of neutrophils is not fully elucidated, as they have been shown to exhibit both protective and harmful capacities in TB. This duality suggests possible differences in the nature and type of neutrophils present during the infection, generating different effects. We hypothesized that Mtb infection induces changes in neutrophil phenotype and function, influencing the infection outcomes. In order to decipher the link between neutrophils and disease progression, we used a cynomolgus macaque model of human TB. Based on clinical, bacteriological, and positron emission tomography with X-ray computed tomography (PET/CT) scan parameters, animals were stratified into two categories: animals that rapidly progressed to an active form of TB, designated as “fast progressors”, and “slow progressors”, which include low symptomatic or asymptomatic animals. In this study, we identified transcriptomic signatures of type I interferons and neutrophil degranulation in macaques with fast progression to active TB, which were not observed in animals with slow TB progression. Unsuppervised mass cytometry analysis showed the emergence of blood immature neutrophils (CD101+ CD10-) in fast progressing animals. In addition, circulating neutrophils from infected animals displayed capacities to modulate TNF-α production and cytotoxic function of CD8 T cells in a contact-dependent mechanism. In the lungs, neutrophils infiltration in granuloma was higher in fast progressors and specifically located in the lymphocyte-rich region in lesions. These data suggest that specific neutrophil’ subpopulations are associated with disease progression. Furthermore, these data suggest that neutrophils may modulate CD8 T cells’ functions, which in turn contribute to the loss of Mtb control and fuel inflammation.

**AUTHORS SUMMARY:** *Mycobacterium tuberculosis* (Mtb) infection in humans is associated with a wide range of disease progression, ranging from latent tuberculosis (TB) to active TB. Understanding immune factors leading to the control of the infection or disease progression is essential to identify new biomarkers and targets for host-directed therapies. Innate immunity plays an important role in inflammatory imbalance observed in active TB, among which neutrophils have both beneficial and detrimental roles. Using a macaque model developing a broad range of clinical forms of TB, we seek to understand the links between neutrophils and disease progression. We found that rapid progression to active TB leads to type I interferon signalling and neutrophil activation. In the blood, immature neutrophils were enriched when the disease progressed. In case of severe TB, Neutrophils also infiltrate a specific region of lung TB lesions rich in T lymphocytes, whereas they could modulate CD8 T cells. Our study provides new insights into the role of neutrophils in TB progression.

## INTRODUCTION

Tuberculosis (TB), caused by *Mycobacterium tuberculosis* (Mtb), remains a leading cause of death worldwide, accounting for 1.3 million deaths in 2023 according to the global TB report of the World Health Organization (WHO)(1). TB is continuing to spread, with about 10.8 million new diagnoses of TB in 2023. *Mtb* infection is associated with clinical diversity and outcomes ranging from latency to active TB. However, our understanding of the immune mechanisms relating to protection against TB disease or its progression is limited. Innate immune responses, especially those involving myeloid cells, are thought to participate in Mtb control, shaping the outcome of infection(2). The role of one particular set of myeloid cells, neutrophils, has yet to be fully elucidated as these cells have been shown to have both protective and harmful properties in TB.

Previous studies have highlighted the importance of neutrophils in Mtb clearance and protective immunity against Mtb infection(3–8). Conversely, high levels of neutrophils in the blood and lung tissue are correlated with TB progression in patients(9–11). In addition, uncontrolled infection results in an increase in neutrophil recruitment to the lungs via the production of inflammatory cytokines and has been associated with an aggravation of the disease, including hyperinflammation and necrosis(12–15). These conflicting dual roles of neutrophils suggest possible differences in the nature and type of neutrophils present in different conditions, generating different effects. Indeed, recent studies in the field of cancer have revealed the presence of both antitumoral and protumoral neutrophils, influencing adaptive responses and angiogenesis(16, 17). A few studies on TB have indicated that certain neutrophil subsets are associated with the severity of TB disease(18). For instance, higher frequencies of banded neutrophils are associated with more severe lung disease in TB patients(9, 11). Recent studies have demonstrated the generation of neutrophil subpopulations in Mtb-infected mice and the ability of these cells to modulate the adaptive immune response or drive immunopathology(18, 19). Doz-Deblauwe et al. described inflammatory MHC-II^−^ PD-L1^lo^ inflammatory neutrophils responsible for exacerbated inflammation in accelerated infection, whereas MHC-II^+^ PD-L1^hi^ neutrophils were associated with T-cell suppression in slow TB progression(18). Saqib et al. identified mature CD101^+^ neutrophils providing protection against severe lung disease, whereas immature CD101^−^ neutrophils and type I IFNs promoted lung damage(19). However, the diversity of neutrophils in TB and its implications remain poorly understood, partly due to the challenges faced in laboratory studies of neutrophils. There is also a lack of consensus concerning the definition of neutrophil subpopulations, as various studies have used different strategies and methods. The neutrophil populations of mice and humans also differ in terms of granule content^20^, cytokine production^21^, phenotype^22^, and response to infection, limiting the translation of findings from mouse models into clinical studies. Non-human primates (NHPs), such as cynomolgus macaques, are a valuable preclinical model of TB, as they reproduce the clinical heterogeneity observed in humans, in animal species with a very similar immune system and respiratory physiology(20, 21). We recently defined maturation-based neutrophil subpopulations in Mauritian cynomolgus macaque combining the expression of CD101 and CD10 with a phenotype and cytological features similar to those observed in humans(22). We used mass cytometry, a high-dimensional single-cell tool, with unsupervised deep profiling to investigate the complexity of neutrophil subsets in a cynomolgus macaque model of human TB. We hypothesized that Mtb infection induces changes in neutrophil phenotype and function, influencing infection outcome, and we investigated the role of neutrophils in TB progression. We performed low-dose exposure (25 CFU) with Mtb Erdman to reproduce the clinical heterogeneity of human TB, with a range of disease progression rates, in Mauritian cynomolgus macaque(23). We classified the animals into two groups on the basis of clinical, bacteriological, and positron emission tomography with X-ray computed tomography (PET/CT) scan parameters: animals displaying rapid progression to an active form of TB (within 11 to 16 weeks post-infection), referred to here as “fast progressors” (FP), and “animals with only mild symptoms or no symptoms at all, with or without occasional positive Mtb culture results, referred to here as “slow progressors” (SP). The rapid progressors displayed an upregulation of pathways associated with type I interferon and neutrophil signaling not observed in slow progressors, together with an increase in the number of immature neutrophil clusters in the blood. In infected animals, circulating neutrophils acquire the ability to modulate CD8^+^ T-cell cytotoxicity and TNF-α production. In lung granulomas, a greater infiltration of neutrophils into lymphocyte-rich regions was observed in FP than in SP. These findings highlight the importance of neutrophils in the control of Mtb infection, inflammation, and disease outcome.

## RESULTS

### Low-dose Mtb infections in cynomolgus macaques result in heterogeneous disease presentations

We investigated early myeloid immune responses during tuberculosis, by infecting 10 cynomolgus macaques by bronchial exposure to 25 CFU *Mycobacterium tuberculosis* Erdman. We performed longitudinal monitoring, including clinical observation, bacteriology, IFN-γ ELISpot and serial positron emission tomography with X-ray computed tomography (PET/CT) scans with 2-deoxy-2-^18^F-deoxyglucose (^18^F-FDG) (Figure 1, Supplementary Figure 1). Consistent with previous reports(23–25), we found that low-dose *Mtb* infection led to different outcomes in cynomolgus macaques, ranging from a rapid progression to active TB with survival below 20 weeks to slow disease progression with few clinical symptoms. Exposure to the bacterium led to Mtb infection in all animals, as demonstrated by positive ESAT-6/CFP-10 IFN-γ ELISpot responses and lung granulomas observed on PET/CT-scans four weeks after exposure (Figure 1A-C, Supplementary Figure 1D-J). Six of the 10 animals displayed rapid progression to TB disease, with moderate to severe symptoms, reaching the human endpoint from 12 to 20 weeks after exposure (Figure 1B). These animals had a significantly higher clinical score than slow progressors, with weight loss, severe dyspnea, hypoxemia and coughing appearing from week 10 to 16 post exposure (p.e.) and worsening over time (Figure 1B, Supplementary Fig 1A-C). Monocytosis was also observed in all six animals at the onset of severe symptoms (Supplementary Fig 1E). By contrast, the other four macaques displayed slower disease progression, with no symptoms or only light symptoms, consisting of weight loss from week 12 p.e. and/or a transient decrease in blood oximetry findings (3 macaques displayed mild symptoms and one was asymptomatic). Mtb was consistently detected in the BAL of all six fast progressors, transiently in one slow progressor and was never detected in the BAL or the remaining three slow progressors (Supplementary Fig. 1F-G). Due to the disparity of disease progression, clinical presentation and of Mtb detection in fluids, with differences in kinetics and magnitude between animals, we separated the animals into two groups: fast progressors (FP, six, all with active TB) and slow progressors (SP, four, three with different stages of subclininal TB — MF1, MF2 and MF3 — and one with latent TB, MF4).

**Figure 1.**
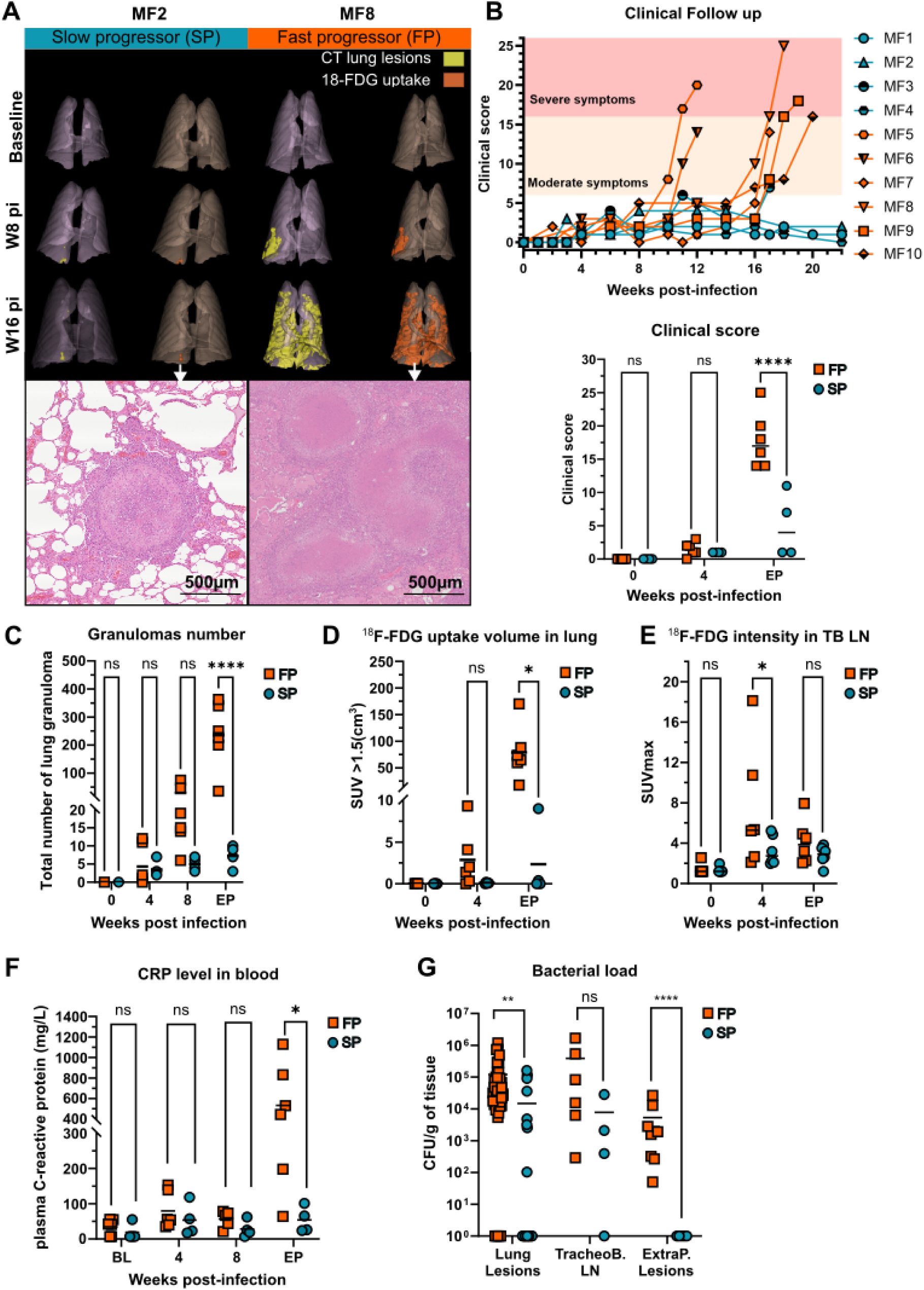
Low-dose Mtb infection leads to heterogeneous clinical progression, lung inflammation and bacterial burden. (A) Three-dimensional reconstructions of the lungs from PET/CT-scans of two representative *Mtb*-infected NHPs at different time points. CT opacities are depicted in yellow and 18-FDG uptake (SUV>1.5) is shown in orange. Representative hematoxylin-eosin staining from caudal right lobe sections of MF2 and MF8. (B) Top panel: Clinical scores for each animal, FP in orange and SP in blue, calculated with the TB-specific scoring grid. A clinical score below 6 is considered asymptomatic, scores between 6 and 16 correspond to moderate symptoms and scores above 16 correspond to severe symptoms. Bottom panel: comparison of clinical scores between FP and SP at baseline, 4 weeks p.e.. Bars indicate the median scores, **** *p*-value< 0.0001, two-way ANOVA with Sidak’s correction for multiple comparisons. (C) Total number of lung granulomas counted on CT-scans. (D-E) ^18^F-FDG uptake volume in the lung with SUV>1.5 corresponding to the FDG-positive threshold, and the ^18^F-FDG signal intensity in tracheobronchial lymph nodes (TB LN), expressed as SUVmax. (F) Plasma C-reactive protein concentration. The data presented are median values, with * *p*-value < 0.05, ** *p*-value < 0.01 and *** *p*-value < 0.001, **** *p*-value< 0.0001, two-way ANOVA with Sidak’s correction for multiple testing. (G) Bacterial load in the lungs and extrapulmonary lesions sampled at necropsy (the following samples were collected: one lesion per lung lobe for each animal, one sample per tracheobronchial lymph node, and one sample from the spleen, liver or kidney). ** *p*<0.01 and \*\*\**p*<10^−3^ in Mann-Whitney *U* tests. BL: baseline, EP: endpoint, SUV: standardized uptake value. Dark cyan: slow progressors (SP) and orange: Fast progressors (FP). LN: lymph nodes.

Three-dimensional reconstructions of serial PET/CT scans highlighted the formation of initial lesions at week 4 p.e. in all animals and revealed differences in lesion progression, including kinetics and dissemination, between animals. Indeed, we observed few isolated hypermetabolic (^18^F-FDG positive) lung granulomas in SP and more diffuse lung lesions with lobular or lobar pneumonia in FP at the latest time point p.e. (Figure 1A, C and D, Supplementary Figure 2). FP had significantly larger numbers of granulomas than SP (*p*<10^−4^) and a greater volume of ^18^F-FDG-positive lung lesions at the time of necropsy (*p*<0.05), highlighting major hypermetabolism. Tracheobronchial lymph node hypermetabolism was observed earlier (Figure 1E), with a standardized uptake maximum value (SUVmax) for ^18^F-FDG at 4 weeks p.e. that was significantly higher for FP than for SP (*p*<0.05). In addition to lung inflammation evidenced by hypermetabolism, the FP displayed significant systemic inflammation with plasma C-reactive protein (CRP) levels at the endpoint higher than those in SP (*p*=0.038), in which these levels remained stable over time (*p*=0.856) (Figure 1F). At necropsy, FP macaques had a significantly higher bacterial load in lung granulomas than SP (*p*<0.05, Figure 1G), whereas the tracheobronchial lymph node bacterial load did not differ between groups (Figure 1G, *p*=0.174). At least one lung granuloma per animal was Mtb-positive in all animals except MF3. However, only FP developed extrapulmonary TB with granulomas in the liver, spleen, and kidneys, highlighting the severity of the infection in these animals (Figure 1G).

Thus, the progression to active TB in FP was associated with greater lung and systemic inflammation, together with severe respiratory symptoms and extrapulmonary TB. By contrast, SP controlled Mtb infection more effectively, displaying less inflammation, with, slowly progressing lung lesions and no dissemination of Mtb to other organs.

### Fast progression of Mtb infection is associated with myeloid cell activation and inflammatory transcriptional signature landscape

For identification of the immune signatures associated with fast or slow disease progression, we performed bulk RNA sequencing on whole blood taken on six macaques at baseline and during the early (D4, W1, W2 p.i), mid (W6, W8 p.i) and late (W12, W14, W16 pi) phases of infection. We identified 855 differentially expressed genes (DEGs), 501 of which were upregulated and 354 downregulated in FP relative to SP (Supplementary Figure 3A). We characterized the pathways associated with disease progression by performing functional gene set enrichment analysis (GSEA) on DEGs identified in comparisons between FP and SP, using Gene Ontology and Reactome (Figure 2A-B). In Gene Ontology analysis, rapidly progressing TB infection was linked to a significant upregulation of biological processes (*p*-adjust<0.05) including innate immune response, defense response to other organisms, bacteria and viruses (Figure 2A). This analysis also revealed a significant upregulation of the cellular component pathways of neutrophil granules in FP: tertiary granules, secretory granules and specific granules. Reactome pathway analysis showed significant enrichment in a neutrophil degranulation biological process, along with γ and α/β interferon signaling, antimicrobial peptides and alpha-defensins in FP relative to SP (Figure 2B). We then focused our analysis on the top-ranking DEGs associated with the Gene Ontology response to bacterium and neutrophil granule pathways (Figure 2C). The defense against bacteria-associated pathway includes interferon-stimulated genes (*GBP2-6, IRF2, JAK2* and *OAS2-3*) and class-I MHC genes (*MICA* and *MR1*) upregulated in FP. Finally, we analyzed the contributions of the most variable DEGs to Gene Ontology pathways (Figure 2D). The dominant genes contributed to innate immune responses and neutrophil degranulation processes (cytoplasmic vesicle lumen, secretory granule lumen, tertiary granule lumen and vesicle lumen). They included genes encoding granule proteins (*MMP8, LCN2, S100A8, DEFA5, S100A9, LTF, HP, CRISP3, FOLR3, TNFAIP6* and *SERPINB1*), a pattern-recognition receptor (*PGLYRP1*) and interferon-stimulated genes (*IFI27, ISG15, IFIT2* and *MX2*) (Figure 2D). Overall, our data indicate that the rapid progression of *Mtb* infection in FP is linked to various transcriptional changes, with a large contribution of neutrophil signatures. Our analysis reveals increases in neutrophil degranulation, and in type I and type II interferon signaling, the latter corresponding to transcriptomic signatures observed in patients with active TB but not in those with latent TB(26).

**Figure 2.**
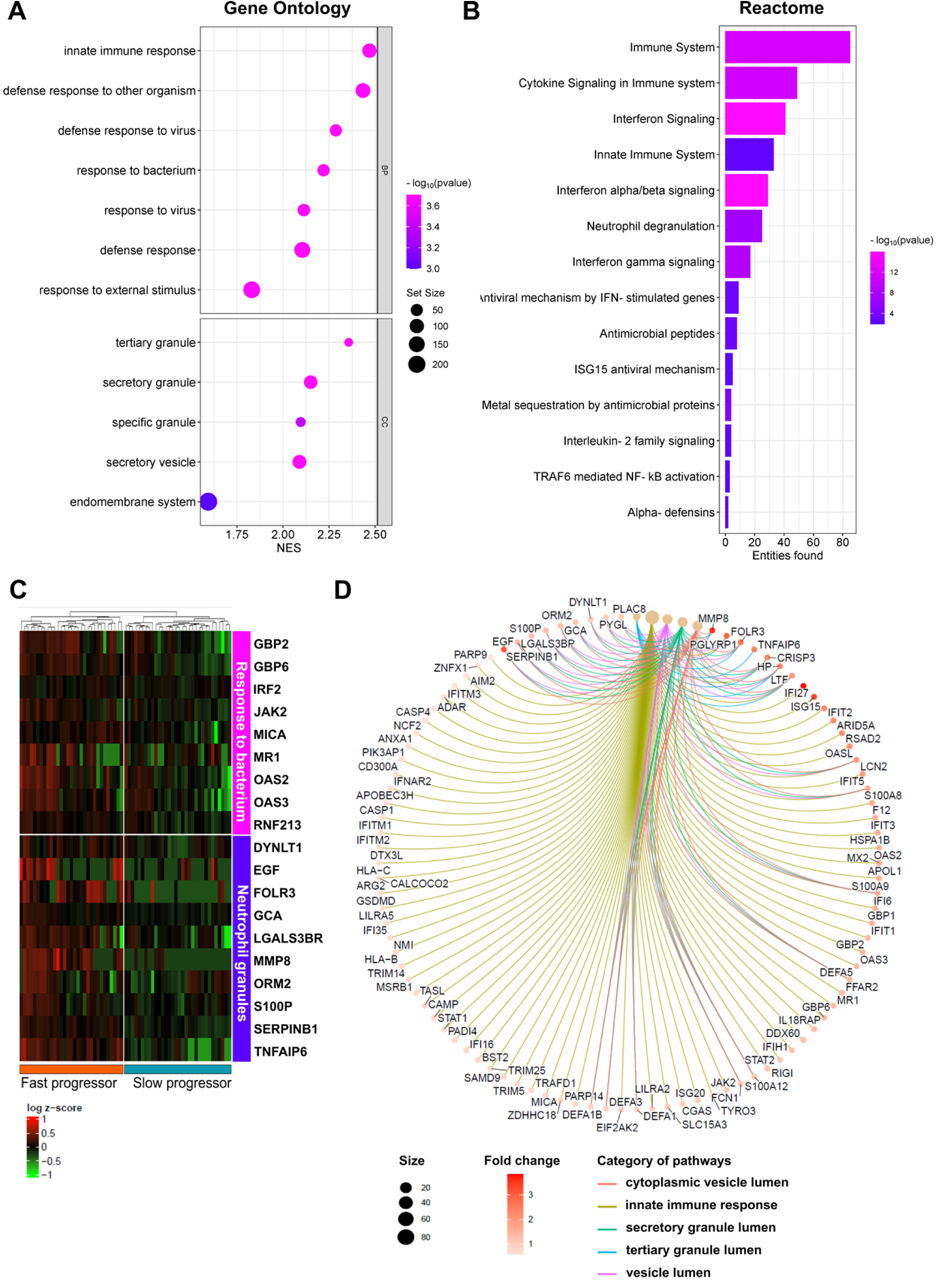
Fast progression of Mtb infection is associated with neutrophil and inflammatory cytokine transcriptomic signatures. (A) Pathways enriched in FP versus SP, according to the DEGs, for Gene Ontology biological processes. The size of the bubble indicates the proportion of genes, and the color indicates the significance (adjusted *P*-value). NES is the enrichment in FP relative to SP. (B) Pathways enriched in FP versus SP, according to the DEGs, in Reactome analysis. Bars indicate the number of genes enriched and the color indicates the significance (adjusted *P*-value). (C) Heatmap of DEG (in rows) associated with Gene Ontology pathways of interest: Response to bacterium and neutrophil granules, differing between SP and FP at late time points analyzed. (D) Connectivity netplot of functional enrichment pathways from GO pathway analysis with the most variable genes. GO: gene ontology, BP: biological processes, DEG: differentially expressed genes, NES: normalized enrichment score.

### Neutrophil subsets display differential maturation profiles during the progression of TB

Transcriptomic analysis showed that pathways associated with neutrophil activation and degranulation were upregulated in FP relative to SP. We investigated myeloid cell responses during *Mtb* infection, by using mass cytometry to characterize the myeloid cells in blood and BAL. The mass cytometry data were subjected to unsupervised analysis by FlowSOM clustering and uniform manifold approximation projection (UMAP) dimensional reduction. The FlowSOM clustering of blood and BAL cells generated 82 and 100 clusters, respectively, each associated with a unique phenotype and abundance trajectory (Figure 3 and Supplementary Figure 4). The clusters were then grouped into 14 metaclusters corresponding to major leukocyte populations from blood and BAL, according to a combination of markers defining immune cells in cynomolgus macaques (Figure 3A and 3B and Supplementary Table 2). Neutrophils were the most diverse population in the blood in terms of the number of clusters, with 32 distinct clusters displaying different phenotypes throughout disease progression and over different time points (Figure 3B-C). Blood monocytes represented 6 cluster in total, showing significant increase at endpoint in FP with minor changes at cluster level (Supplementary Figure 4B-E). There were only 10 neutrophil clusters in BAL, whereas macrophages displayed greater heterogeneity and abundance, with 45 clusters in total (Figure 4C and Supplementary Figure 4F-H).

**Figure 3.**
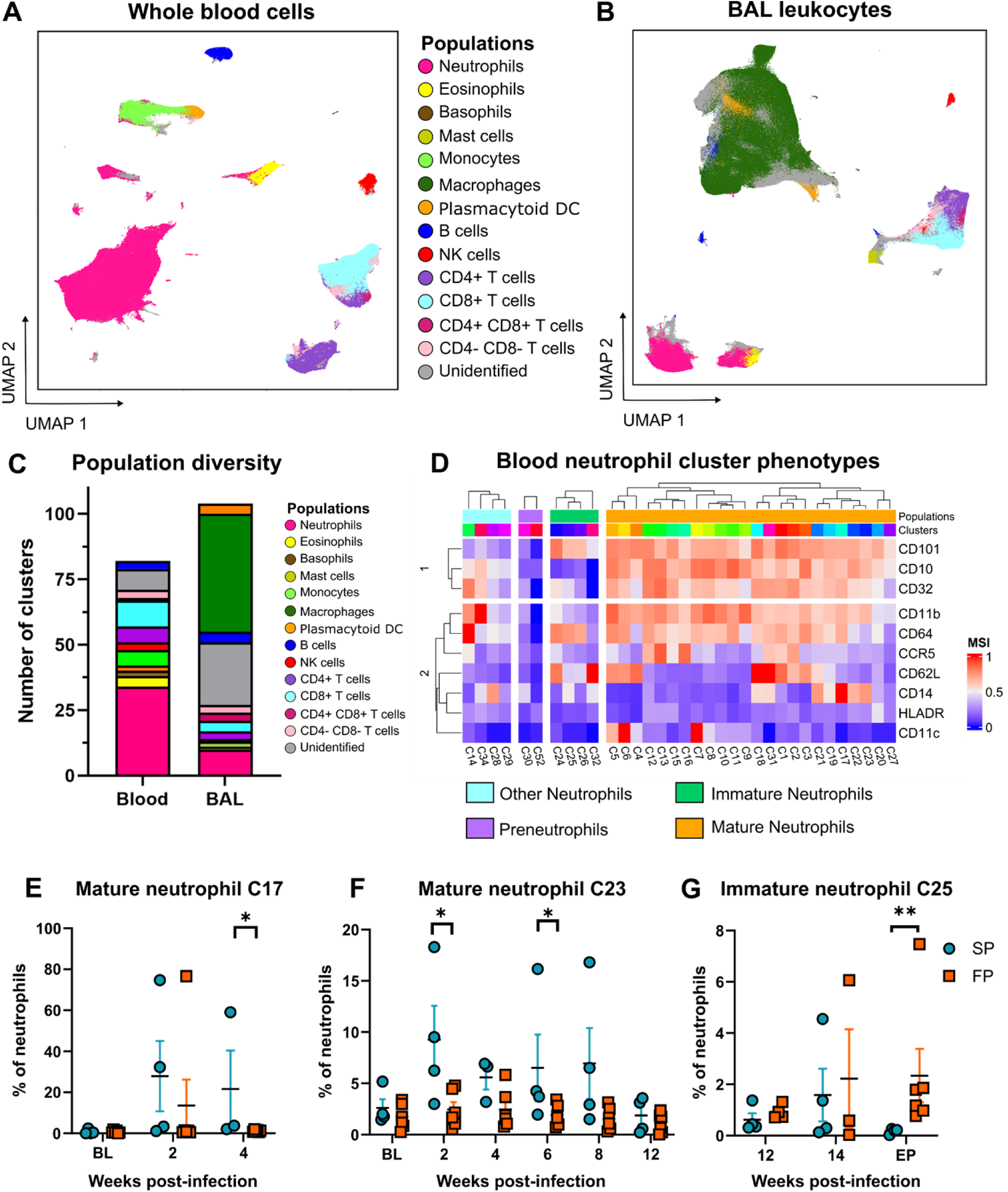
Mtb infection and progression is associated with a shift in neutrophil maturation. (A-B) UMAP visualization of CyTOF whole blood cell populations and BAL leukocytes, colored by generic cell type: “metaclusters”. (C) Population diversity, represented by the number of unique clusters identified with FlowSOM in blood and BAL. Stack bar plot colored by cell population. (D) Heatmap showing phenotypic clustering of whole-blood neutrophils. Clusters are spread in columns and grouped into subpopulations of neutrophils. The expression level of each surface marker is depicted in rows, the color indicating the intensity of the signal of each marker from low (blue) to high (red) intensity. (E-G) Comparison of the percent abundance of whole-blood neutrophils clusters between SP and FP at different time points. MSI: 1: high levels of expression and 0: no expression. is the data are presented as the mean ± standard mean of error (SEM), with * *p*-value < 0.05, ** *p*-value < 0.01, for Mann-Whitney *U* tests at the time points indicated.

**Figure 4.**
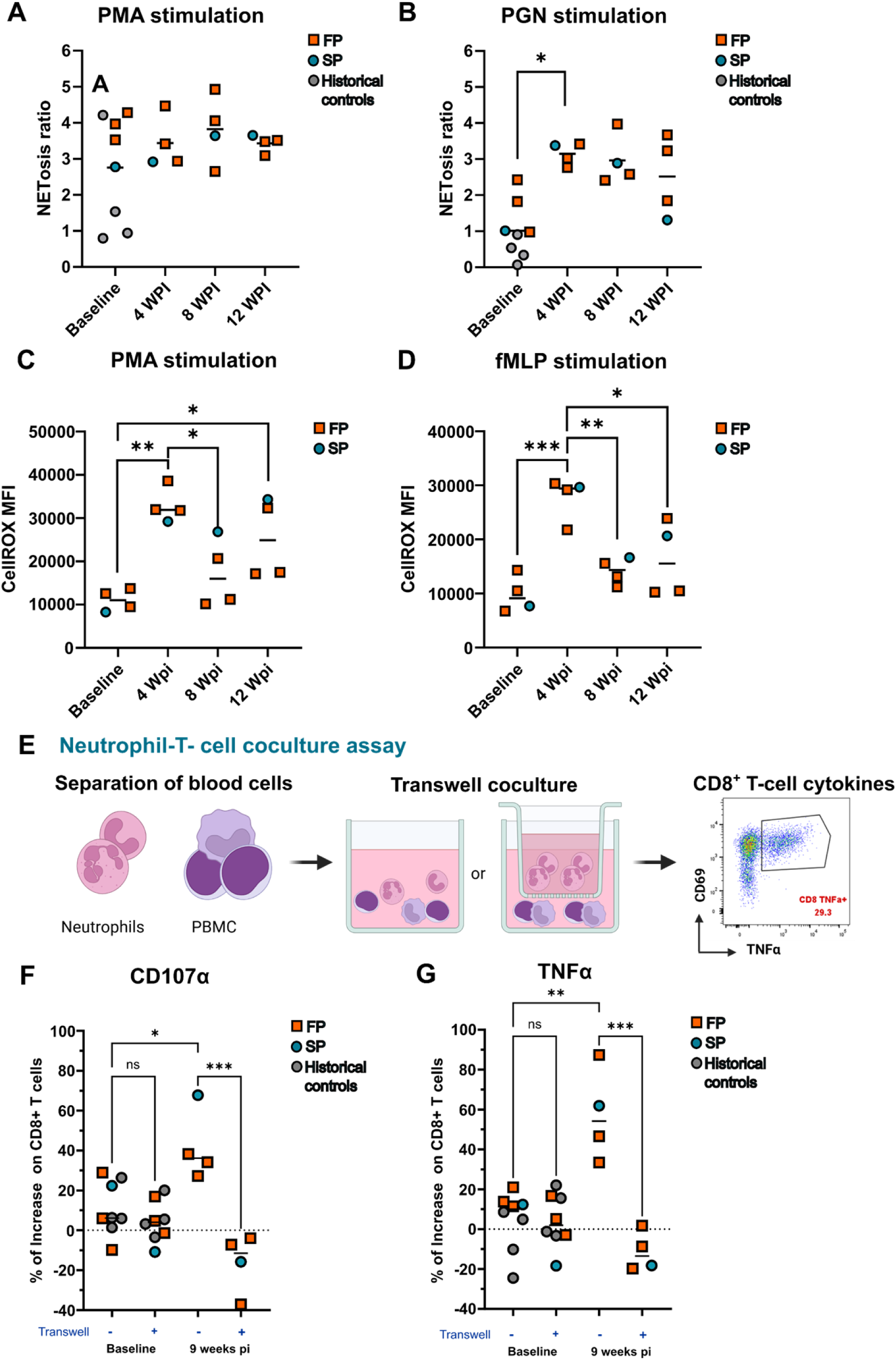
Functional characterization of neutrophils. (A-B) NETosis ratio of blood neutrophils stimulated with phorbol myristate acetate (PMA) or peptidoglycan, quantified by MPO-DNA ELISA at different time points. The horizontal bar represents the mean MFI at each time point. \**p*<0.05 in a Kruskal-Wallis test with Tukey’s correction. (C-D) Reactive oxygen species (ROS) production by whole-blood neutrophils stimulated with PMA or N-formyl-L-methionyl-L-leucyl-L-phenylalanine (fMLP), measured by flow cytometry with a CellROX probe. The signal is expressed as the mean fluorescence intensity of the ROS marker (CellROX) and the horizontal bar represents the mean MFI. \**p*<0.05, \*\**p*<0.01, \*\*\**p*<0.001 in a Kruskal-Wallis test with Tukey’s correction. (E) Diagram of the design of the neutrophil-T cell coculture and Transwell assay. Sorted neutrophils were cocultured with autologous PBMC in the presence or absence of a Transwell membrane. Activation markers and cytokine production by T cells were measured by intracellular staining coupled with flow cytometry after overnight stimulation with Cytostim. BioRender.com. (F-G) Percent increase in CD107a expression and TNF-α production by CD8^+^ T cells correspond to the increase or decrease in the percentage of CD107a^+^ or TNFα^+^ cells in the presence of neutrophils relative to PBMC alone. Data collected from animals nine weeks after infection were compared with baseline and historical controls, i.e. healthy animals from other internal studies. \**p*<0.05, \*\**p*<0.01, \*\*\**p*<0.001 iny Kruskal-Wallis tests with Tukey’s correction.

Before infection, we observed no major differences between the two groups for any of the clusters on UMAP observation and abundance analysis (Supplementary Figure 4A). However, using opti-SNE and focusing on neutrophils, we observed that Mtb infection was associated with changes in blood neutrophil cluster abundance in both SP and FP (Supplementary Figure 5A and 5B). Blood neutrophil clusters were grouped together according to the expression of CD10, CD101 and CD32, delineating clusters corresponding to preneutrophils (CD101^−^ CD10^−^ CD32^−^), immature neutrophils (CD101^+^ CD10^−^ CD32^−/low^) and mature neutrophils (CD101^+^ CD10^+^ CD32 ^low/high^) (Figure 3D, Supplementary Figure 5D), as in a previous study(27). The unsupervised analysis also revealed another population of CD101^−^ CD10^low/mid^ CD32^−/+^ neutrophils (referred to as “other neutrophils”), which we did not observe in previous studies on cynomolgus macaques. In SP, at week 12 and 14 p.e., and at the endpoint, there was a shift of mature CD101^+^ CD10^+^ neutrophils towards CD62L^low^ activated phenotypic clusters (C10, C7 and C27) (Supplementary Figure 5B). In FP, at early (2 wpi and 4 wpi) and intermediate time points (6 wpi to 12 wpi), there was a decrease in the abundances of mature CD101^+^ CD10^+^ CCR5^+^ clusters (C2, C13, C15 and C16), whereas at later time points (14 wpi to 18 wpi), an enrichment in mature activated CD62L^low/mid^ CCR5^−^ clusters (C5, C6, C8 and C10) was observed. FP had a greater abundance of mature neutrophil clusters (C23 and C17) than SP early in Mtb infection, especially at week 2 and 4 p.e. (*p*=0.0238, *p*=0.0381, respectively) (Figure 3E and 3F). Cluster 23 remained significantly more abundant at 6 weeks pi (p=0.0381, Figure 3F). Phenotypically, C17 and C23 were activated CD11b^high^ CD62L^low^ cells also expressing CD14 but differing in CD64 expression, which was observed only in C17, as depicted on the phenotypic heatmap (Figure 3D). Late in infection, the immature neutrophil cluster C25, characterized as CD11b^low^CD64+CD14^mid^CD62L^low^ cells, was significantly more abundant in FP than in SP (*p*=0.00952, Figure 3D and 3G, Supplementary Figure 5B). We then grouped the clusters into mature, immature, preneutrophil and other neutrophil subpopulations, but we found no significant difference in the abundance of these subpopulations between groups (Supplementary Figure 5D).

In BAL, no significant differences in the frequencies of neutrophil clusters were observed between the two groups (Supplementary Figure 5C), whereas two clusters of alveolar macrophages (C30 and C22) were significantly more abundant in FP than in SP (Supplementary Figure 4G-H). Overall, the rapid progression of Mtb infection was associated with early changes in neutrophil subpopulations detected only in blood. In particular, immature neutrophil clusters progressively increased in abundance with disease progression.

### Progression of Mtb infection is associated with NETosis and a modulation of the-cell response by neutrophils

We investigated the possible association of changes in neutrophil maturation with modulation of the functions of these cells, by assessing NETosis and ROS production in four cynomolgus macaques infected with Mtb (three FP and one SP). NET production by blood neutrophils was quantified by determining the amount of myeloperoxidase-DNA (MPO-DNA) complex produced after stimulation with peptidoglycan (PGN) or phorbol myristate acetate (PMA) (Supplementary Figure 6). With this system, unstimulated neutrophils displayed negligible or absent NET release, whereas PMA stimulation induced the release of large numbers of NET, with no significant differences before and after infection (Figure 4A and 4B). We also used PGN as a physiological stimulus inducing NET production in neutrophils. NETosis levels were significantly higher than those in uninfected macaques 4 weeks p.e. (*p*<0.05), indicating that the infection induced an increase in the capacity of neutrophils to release NET. NET production was not significantly different at later time points, although the data were more heterogeneous. We also determined reactive oxygen species (ROS) in a whole-blood flow cytometry assay based on cellROX technology. The capacity of neutrophils to produce ROS in response to PMA or N-formyl-L-methionyl-L-leucyl-L-phenylalanine (fMLP) was found to have increased significantly 4 weeks p.e. (*p*<0.01 and *p*<10^−3^, respectively) (Figure 4C and 4D). ROS production levels in response to stimulation with PMA or fMLP were significantly lower at 8 weeks p.e. than at 4 weeks p.e. (*p*<0.05 and *p*<0.01, respectively). At a later time point (12 weeks p.e.), the ROS response of neutrophils to PMA remained significantly stronger than that at baseline. These data indicate that the capacity of neutrophils to produce NET and ROS increases early in infection. At 4 weeks p.e., only neutrophil cluster 17 displayed a decrease in frequency. It remains to be determined whether these functions change during disease progression.

In other studies, changes in neutrophil phenotype or neutrophil subsets have sometimes been shown to be associated with capacities to modulate T cells(18). We previously showed that neutrophils could modulate T cells during acute and chronic simian immunodeficiency virus infection in cynomolgus macaques (Lemaitre et al., 2022). We therefore assessed the capacities of purified autologous neutrophils to modulate CD4^+^ and CD8^+^ T-cell activation, cytotoxicity and intracellular cytokine production in an *in vitro* coculture assay (Figure 4E, Supplementary Figure 7). Due to the large volumes of blood required, we limited our analysis to two time points: baseline and 9 weeks p.e. in four cynomolgus macaques infected with Mtb. Before infection and in historical uninfected controls, neutrophils had a limited impact on T-cell functions. At 9 weeks post-infection, the neutrophils from infected animals had increased the levels of CD107a expression and TNF-α production by autologous CD8^+^ T cells relative to baseline (Figure 4F and 4G). We investigated the mechanism of T-cell modulation by neutrophils in a Transwell assay in which neutrophils were separated from PBMC in two medium-filled compartments separated by a membrane with 0.4 µm pores. The presence of this membrane decreased T-cell modulation by neutrophils, highlighting the existence of a contact-dependent mechanism (Figure 4F and 4G). These findings indicate that neutrophils upregulate the cytotoxicity and TNF-α production of T cells during Mtb infection, via a contact-dependent mechanism. This mechanism seems to be independent of disease progression status at the time point considered, or there may have been too few individuals to distinguish differences between SP and FP.

### Neutrophil infiltration is associated with disease progression and a specific localization within the lung granuloma

Transcriptomic and mass cytometry analyses highlighted a major link between disease progression and neutrophil activation, degranulation, and differentiation. We investigated the possible association between disease progression and a specific pattern of neutrophil distribution within lung granulomas by performing immunohistochemistry for calprotectin on caudal and cranial lung lobe sections (Figure 5A and 5B). We used QuPath software to define four regions of interest (ROI) for the measurement of neutrophil infiltration in the lung: normal parenchyma, lesional parenchyma, necrosis transition, macrophage cuff, and lymphocyte cuff. Rapid progression of the infection was accompanied by a significant increase in neutrophil infiltration in the lung (*p*<0.05) (Figure 5C). Analysis restricted to each of the four ROIs showed neutrophil enrichment only in the lesional parenchyma (*p*<0.01) and lymphocyte cuff (*p*<0.01) (Figure 5D-G). These results indicate a preferential infiltration of neutrophils into T cell-rich areas in FP macaques, suggesting strong interactions between these two immune cell populations.

**Figure 5.**
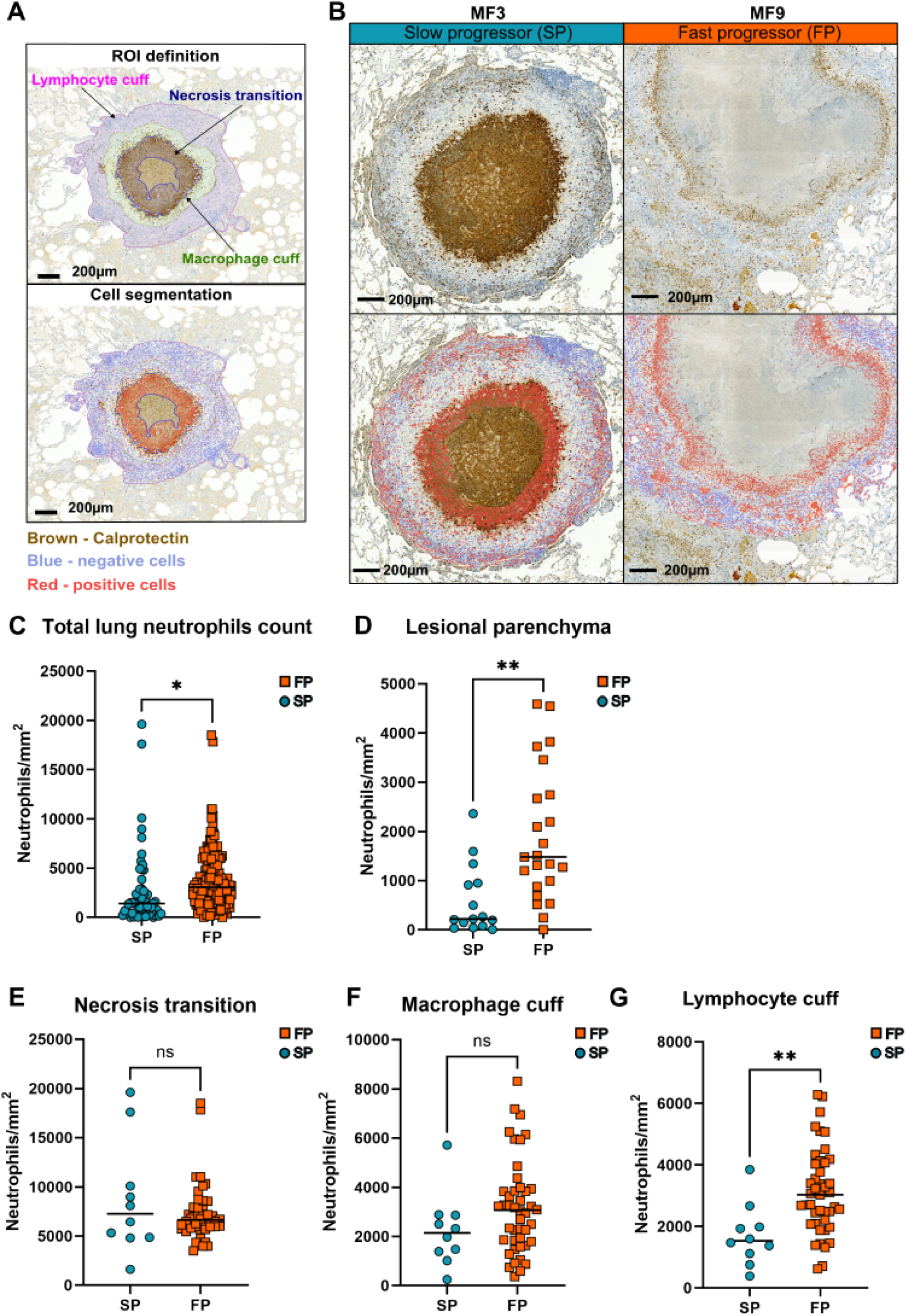
Lung neutrophil infiltration at specific sites in the granuloma is associated with disease progression. (A) The top part of the figure shows the three regions of interest delimited within lung granulomas: necrosis transition, macrophage cuff and lymphocyte cuff. Calprotectin staining appears in brown (DAB). The bottom part of the figure is a representative image from a lung granuloma used for cell segmentation with QuPath. Cells positive for calprotectin staining are depicted in red and cells negative for calprotectin are shown in blue. (B) The top part of the figure showsrepresentative images of calprotectin staining (brown) in caudal left lobes from MF3 (SP) and MF9 (FP). The bottom part of the figure shows the associated cell segmentation on the same images with QuPath, with calprotectin-positive cells in red and calprotectin-negative cells in blue. (C) Total neutrophil count (DAB-positive cells) in all the ROI analyzed. (D-F) Neutrophil counts (DAB-positive cells) in the regions of interest indicated. SP: Slow progressor, FP: Fast progressor. * *p*<0.5, ** *p*<0.01 in a Mann-Whitney *U* test.

### Lung granuloma neutrophil diversity is linked to the blood and spleen compartments

During differentiation and migration to tissues, neutrophils may acquire a tissue-specific phenotype, particularly under inflammatory conditions. We characterized the phenotypic diversity of neutrophils in different organs, to identify tissue-specific phenotypes and potential similarities between the compartments studied. We used mass cytometry and an unsupervised flowSOM approach to analyze and compare neutrophil phenotypes in bone marrow (BM), blood, spleen, tracheobronchial lymph nodes and lung granulomas. Spleen, tracheobronchial lymph nodes and lung granulomas were sampled at necropsy, and BM was obtained at baseline, 4 and 8 weeks p.e. and at necropsy (12 weeks p.e.) from four Mtb-infected macaques (3 FP and 1 SP). Lung granulomas were selected on the basis of PET/CT images, for the analysis of granulomas formed early (4 weeks p.e.) or later (>10 weeks p.e.) during the course of infection, as the structures of these two types of granuloma have been shown to be different(28). We analyzed the spleen, as this organ serves as a reservoir of neutrophils capable of performing extramedullary granulopoiesis and because we observed a few granulomas in the spleen of FP (but never in the spleen of SP). We focused our tissue analysis on infected tracheobronchial lymph nodes (found in all animals), into which neutrophils could infiltrate due to granuloma formation. FlowSOM clustering analysis revealed that neutrophils accounted for 17.0% of lung granuloma cells, 26.83% of spleen CD45^+^ cells and 77.93% of BM cells, but that neutrophils were absent from tracheobronchial lymph nodes (Supplementary Figure 8A and 8B). Using a clustering approach similar to that used for blood cells, we identified 30, 50 and 100 neutrophil clusters in lung granulomas, the spleen and BM, respectively. Based on the expression of CD101 and CD10, we grouped lung, spleen and BM neutrophil clusters into mature neutrophils (CD101^+^ CD10^+^), immature neutrophils (CD101^+^ CD10^−^), preneutrophils (CD101^−^ CD10^−^), and other neutrophils (CD101^−^ CD10^+^) (Supplementary Figure 9A-C). This analysis revealed differences in the frequencies of CD4^+^ T cells, CD8^+^ T cells and macrophages between the granulomas that formed early and those that formed late, but there were too few samples to draw any firm conclusions about the significance of differences (Supplementary Figure 8C and 8D). We therefore pooled the early and late granulomas together for the rest of the analysis. As expected, about 80% of whole-blood neutrophils had a mature phenotype, a percentage significantly higher than that for the spleen, BM and lung granulomas (*p*<0.05, *p*<10^−3^ and *p*<10^−4^) (Figure 6A). The frequency of preneutrophils in the spleen was significantly higher than that in whole blood (*p*<0.05). We observed no significant difference in the frequency of immature neutrophils between compartments on necropsy. The presence of preneutrophils and immature neutrophils in the spleen is consistent with previous reports showing that the spleen is capable of extramedullary granulopoiesis(29). Finally, the frequency of other neutrophils was significantly higher in lung granulomas than in whole blood, bone marrow and spleen (*p*<0.05), suggesting the existence of a lung granuloma-specific phenotype. An analysis of BM neutrophil phenotype during the course of Mtb infection revealed no major changes at cluster level but highlighted a significant increase in preneutrophil levels between baseline and 12 weeks p.e. (Supplementary Figure 9D-E). We investigated neutrophil diversity within tissues further, with the aim of identifying tissue-related phenotypes, by comparing and linking clusters between compartments with the Cytocompare algorithm(30) (Figure 6B-D, Supplementary Figure 9F and 9G). Based on the expression of 10 neutrophil markers (Supplementary Figure 9A-C), we were able to identify phenotypically similar clusters of mature neutrophils (Figure 6C), immature neutrophils, preneutrophils, and other neutrophils (Figure 6D) in the BM, blood, lung granulomas, and spleen. Interestingly, the neutrophil clusters of the spleen and lung granulomas were the most similar, with a total of 61 phenotypic links between these two compartments, probably because granulomas were present in both (Figure 6B). Mature neutrophil clusters made the most important contribution to these links, accounting for 31 links in total. By contrast, we only observed 29 links between blood and lung granulomas and 21 links between BM and lung granulomas. Interestingly, the diverse “other neutrophils” clusters was present in both the spleen and lung granulomas (Figure 6D). These results indicate that most lung granuloma neutrophil clusters are enriched in “other neutrophils”. Globally, lung granuloma neutrophils have a phenotype similar to that of spleen neutrophils, suggesting similarities between the granulomas in these two compartments.

**Figure 6.**
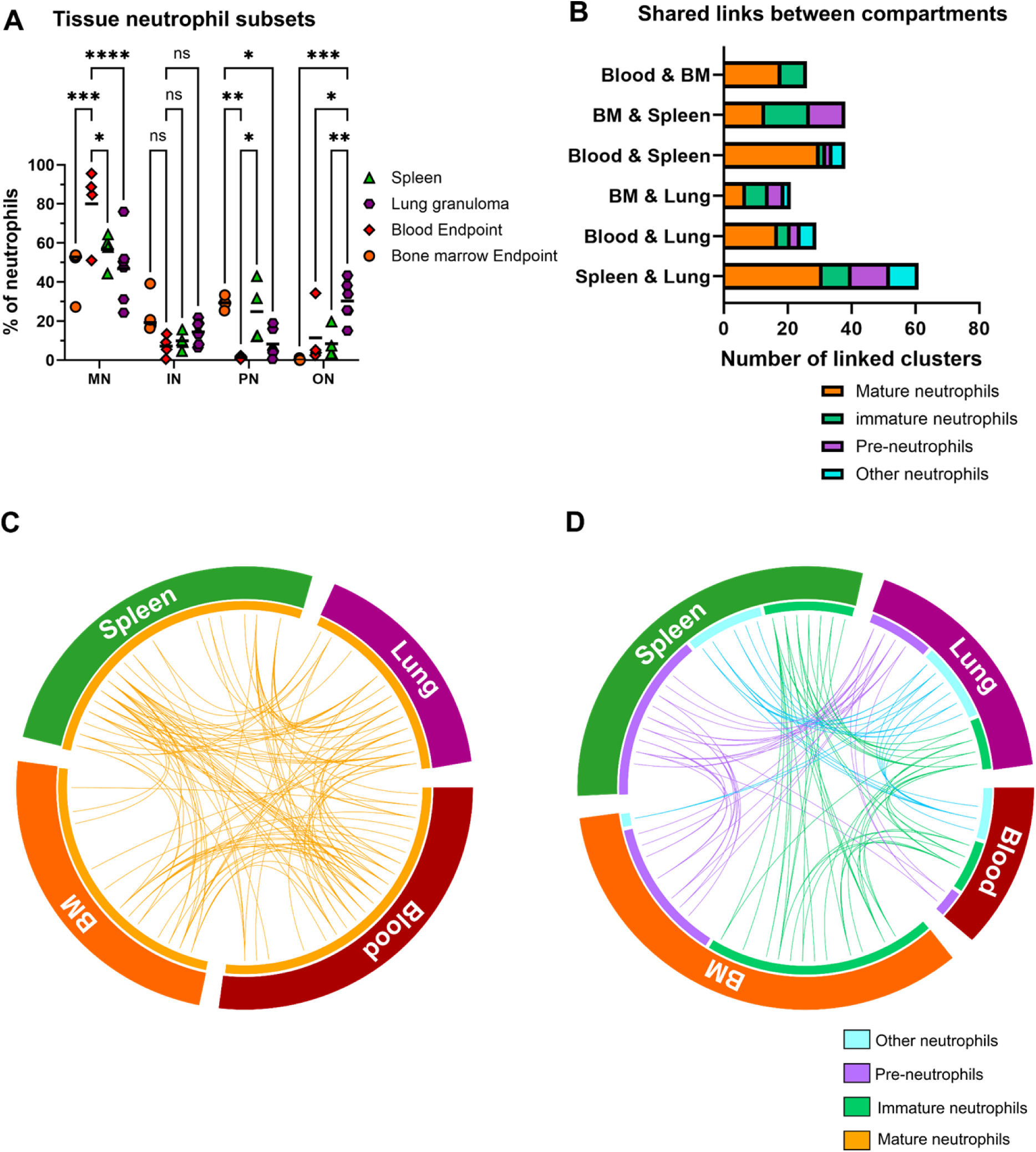
Lung granuloma neutrophil diversity is linked to the spleen compartment. (A) Comparison of the frequency of neutrophil subsets between bone marrow, blood, lung granulomas and spleen at the time of necropsy. The data presented are median values, with * *p*-value < 0.05, ** *p*-value < 0.01 and *** *p*-value < 0.001, *** *p*-value< 0.0001, two-way ANOVA test with Sidak’s correction for multiple testing. (B) Cytocompare analysis comparing and matching neutrophil clusters with similar phenotypes between tissues: spleen, blood, lung granulomas and bone marrow. The stack bar plot represents the number of neutrophils clusters with a similar phenotype in the two tissues. Clusters are grouped together by neutrophil subpopulation. (C-D) Circos diagram showing neutrophil cluster phenotypic connections between bone marrow, blood, lung granulomas and spleen. Neutrophil clusters are colored by subset: preneutrophils, mature, immature and other neutrophils.

## Discussion

Infection with Mtb can lead to a diverse range of disease, known as “the spectrum of TB”. There is no real consensus on the exact definition of each disease state, but several studies have described similar states, including clearance (Mtb eliminated by the innate or adaptive immune response), latency (Mtb infection is controlled, no symptoms and no culturable Mtb), incipient and subliclinical TB (Mtb infection is not fully controlled. It does not cause clinical TB-related symptoms but there are detectable manifestations indicative of disease and Mtb can be detected by radiological or microbiological assays) and active TB (Mtb infection is not controlled and is accompanied by symptoms of various intensities) (31).The identification and precise characterization of these different TB states may make it possible to provide help earlier and to achieve more personalized care for patients, especially given the large potential for transmission worldwide. Such work would also support research and the development of therapeutic and prophylactic strategies. Uncovering the key factors influencing disease progression or regression is fundamental. We know that Mtb lineages responsible for infection have an impact on infection outcome and extrapulmonary dissemination in patients(32), but host factors are also important to explain such clinical diversity(33). The innate immune response is a major host factor contributing to Mtb control or progression, and to lung immunopathology. Neutrophil signatures have been associated with type I and type II interferon signaling in active TB(19, 26, 34). Depending on the study, neutrophils have been shown to be either harmful or beneficial, and these dual but opposite roles suggest that the type of neutrophils recruited during infection may determine the outcome of infection. Here, we characterized the heterogeneity, dynamics, and functions of neutrophils as a function of TB disease progression. We infected 10 Mauritian cynomolgus macaques with low doses of the Erdman strain to model the disease heterogeneity observed in humans(20, 23). We followed and classified disease progression in each animal by 18-FDG PET-CT imaging, clinical monitoring, and the determination of bacterial load in BAL. As previously described, the clinical course of the disease was heterogeneous. We grouped the animals into two categories — “slow progressors” and “fast progressors” — according to their clinical score, PET-CT images and the presence of Mtb in BAL samples. Fast progressors had more lung lesions, greater lung inflammation, and a higher bacterial load in tissues (including extrapulmonary lesions), corresponding to active TB. Conversely, slow progressors had fewer lung lesions, no extrapulmonary TB and were asymptomatic or had only mild symptoms, with Mtb detected only transiently in BAL cultures. Slow progressors matched the definitions of subclinical TB. One of the individuals had no symptoms, no culturable Mtb in BAL and no progression of the lesions that appeared in W4 p.e. during the course of the study. This individual had disease potentially corresponding to latent TB, but the correct validation of latent TB in NHP remains a challenge, requiring longer follow-up (up to 6 months with the features observed). We therefore grouped this animal with the other macaques displaying slow progression. Mauritian cynomolgus macaques infected with a low dose of Mtb Erdman are an interesting model for studying the link between clinical heterogeneity and host-pathogen interactions. Our results for this model demonstrate poorer control of Mtb replication and dissemination, and a stronger systemic and lung inflammatory response in FP, with rapid progression to active TB.

We investigated the immune parameters associated with disease progression by performing blood transcriptomic analysis in Mtb-infected macaques. We detected an upregulation of genes associated with neutrophil degranulation, type I and type II interferon signatures in fast progressors relative to slow progressors. This finding is consistent with previous reports of an overactivation of IFN-γ and type I IFNs pathways in whole-blood neutrophils from patients with active TB(26). In a recent study, transcriptomic analysis on whole-blood sorted neutrophils from patients with active TB also showed an upregulation of neutrophil migration chemotaxis-related genes (*CXCR4* and *CXCL8*), suggesting a mobilization of bone marrow neutrophils(35). However, in the same study, Geng X. et al. showed that neutrophil degranulation was downregulated in patients with active TB(35). This discrepancy might be due to the differences in blood preparation, with neutrophils being sorted from whole blood collected in EDTA tubes. By contrast, we performed whole-blood analysis on cells preserved in Tempus medium. We observed an increase in the persistent activation and degranulation of neutrophils during infection. The increase in degranulation may not only enhance the inflammatory landscape, but may also damage the tissues, through a release of granular proteins associated with NETs. Furthermore, the genes making the largest contribution to the neutrophil degranulation signature were *MMP-8, DEFA, TNFAIP6, EGF, PGLYRP1, FOLR3*, and *S100*. Interestingly, a human TB study showed that MMP-8 expression increases in neutrophils, driving AMPK-dependent tissue destruction, in human pulmonary TB(36). Gopal et al. demonstrated that neutrophils producing S100 proteins are dominant within the inflammatory lung granulomas in active TB in humans and non-human primates(13). They also showed that S100A8/A9 proteins promote neutrophilic inflammation and worsen lung disease in a mouse model. In addition, higher concentrations of alpha-defensins were observed in bronchoalveolar lavage fluid (BALF) in TB patients with cavitary lesions, and the concentrations of these molecules were positively correlated with that of IL-8, a neutrophil chemoattractant(37). TNFAIP6, an inflammation-associated protein, was also shown to be upregulated in active TB in two cohorts from South Africa and Malawi and it was possible to distinguish between ATB from LTB based on the levels of this protein(38). Increases in degranulation may therefore increase inflammation, worsening tissue damage and facilitating disease progression. It remains uncledar whether degranulation pathways are linked to increases in NETosis in tissues because granules play a major role in the NET formation process. Our results show that neutrophil-associated parameters and interferon signaling are strongly associated with disease progression.

We also characterized the dynamics and phenotypic diversity of neutrophils during Mtb infection. The phenotypic diversity of neutrophils has been little studied in TB, whereas it has been associated with a significant impact on pathophysiology in other infectious diseases, autoimmune diseases, and cancers(16, 39). Studies of neutrophil heterogeneity have used different definitions of neutrophil subpopulations based on phenotype, function, cytology, or density gradient-related properties. Some studies have focused on low-density neutrophils, whereas others have focused on activated neutrophils, or immature neutrophils. Mohd Saqib et al. recently showed that the type I interferon response drives CD101^−^ neutrophil recruitment in the lung and is associated with disease progression in mice(19). In mice, CD101 expression can be used to distinguish between mature and immature neutrophils, whereas, in humans and cynomolgus macaques, CD10 is a better marker for distinguishing between mature and immature neutrophils(22, 40). We used mass cytometry to combine 42 markers to obtain a global vision of immune cells and a more precise description of neutrophil subpopulations. We performed mass cytometry staining on whole blood and BAL within two hours of sampling, to avoid neutrophil activation. We previously showed that the combination of CD10, CD101, and CD32a could be used to distinguish preneutrophils from immature and mature neutrophils in cynomolgus macaques(22), as in humans(40). Blood analysis revealed a diversity of neutrophils, with 32 clusters of mature neutrophils (CD10^+^ CD101^+^), immature neutrophils (CD10^−^ CD101^+^), and preneutrophils (CD10^−^ CD101^−^). We also observed a new population of CD101^−^ CD10^+/int^ neutrophils that we refer to here as “other neutrophils”, which was barely detectable in BM. Neutrophils were activated early in the course of Mtb infection, with an increase in activated mature neutrophil clusters as early as 2 or 4 weeks p.e. in FP relative to SP. With progression of the disease to active TB, the neutrophils in the bloodstream switched to an activated immature phenotype at the endpoint, whereas the frequency of these clusters remained low in SP. The abundance of banded neutrophils, corresponding to our immature neutrophils, has been reported to be correlated with the severity of lung disease in humans(9, 11). Immature neutrophils may simply be a marker of disease severity or they may play an active role in lung immunopathology. We therefore hypothesize that infection triggers neutrophil recruitment, triggering signaling to the bone marrow or spleen to generate more neutrophils or new neutrophil subpopulations with specific functional capacities. Mohd S. et al. demonstrated that lung immature neutrophils were mobilized from bone marrow and were responsible for the exacerbation of immunopathology in a mouse model(19). We investigated neutrophil phenotype and infiltration in tissues in more detail by performing mass cytometry on bone marrow, spleen, tracheobronchial lymph nodes, and lung granulomas from four macaques. We observed a significant expansion of the preneutrophil population at 12 weeks p.e.., suggesting a global bone marrow response to the infection. We detected no neutrophils in tracheobronchial lymph nodes, even if a granuloma was present in samples analyzed, suggesting a different cell composition in this type of granuloma. We found preneutrophils, immature neutrophils, other neutrophils and mature neutrophil phenotypes in lung granulomas and the spleen of infected animals, consistent with the presence of granulomas in both the lung and spleen. The neutrophil clusters found in lung granulomas were more similar to those found in the spleen than those found in the blood.

In histopathological analyses, there was more neutrophil infiltration in the lung in FP than in SP, consistent with findings for people with active TB(13). Using cell segmentation and the definition of specific regions of interest, we showed that, in our model, neutrophils preferentially infiltrate lymphocyte-rich areas from the granulomas of FP. The colocalization of these cells raised the question of the capacities of neutrophils to modulate T cells in TB. We found that neutrophils modulated CD8^+^ T-cell activation and functionality, through increases in the expression of TNF-α and CD107a in particular, via a contact-dependent mechanism during Mtb infection. Neutrophils may stimulate the inflammatory, antimicrobial, and cytotoxic activities of CD8^+^ T cells through direct contact in response to Mtb infection(41, 42). Excess CD8^+^ T-cell activation by neutrophils may be one of the mechanisms increasing overall lung inflammation and contributing to disease progression. These findings pave the way for further investigations into the types of effectors involved in this contact-dependent mechanism and the impact of such interactions on infection.

Neutrophil extracellular trap formation, also known as NETosis, is another major mechanism of action of neutrophils against pathogens. Chowdhury et al. recently demonstrated that NETs release is induced by Mtb and promoted by type I interferon(34). NETs were unable to kill Mtb and even promoted its replication, and they also favored necrosis and caseation. Several studies have suggested that the type I interferon associated with NET promotes lung tissue damage, causing necrosis and inflammation(15, 43, 44). We observed an increase in NET production by blood neutrophils following PGN stimulation, which was detectable at 4 weeks p.e. Unfortunately, due to technical issues, we were unable to evaluate NET production following stimulation with Mtb. Another limitation of our analysis is the small number of animals analyzed. We need to improve our understanding of the composition of the NET produced by neutrophils and its impact on other immune cells in the lung, as NET may modulate CD8^+^ T-cell functions within the granuloma.

This study identifies neutrophil subpopulations as important effectors of disease progression or infection control. Our findings link disease progression to type I interferon signaling, neutrophil degranulation, immature neutrophils, and high levels of lung inflammation. Active TB is associated with higher levels of NET production and strong immunomodulation of CD8^+^ T cells by neutrophils. This highlights possibilities for the therapeutic use of drugs to modulate neutrophil functions and/or neutrophil interactions with adaptive immune cells as a means of restoring efficient anti-TB responses. The development of such host-directed therapies will require improvements in our understanding of neutrophil biology in TB and the impact of antibiotic treatments on granulocytes, to make it possible to target the mechanisms underlying this disease in a specific manner.

## MATERIALS AND METHODS

### Ethics statement

Ten cynomolgus macaques (*Macaca fascicularis*) were imported from Mauritius and housed under BSL3 containment at the IDMIT animal facility of the CEA, Fontenay-aux-Roses, France. All animals were confirmed negative for *Mycobacterium tuberculosis* and *bovis* infection before entering the study, based on Primagam, ELISpot, and chest CT scans. The study was conducted in accordance with French national regulations under the supervision of national veterinary inspectors (National authorization number D92-032-02). During the study, IDMIT complied with the standards for Human Care and Use of Laboratory Animals of the Office for Laboratory Animal Welfare (OLAW) under Assurance Numbers #A5826-01 and F20-00448. All experimental procedures were conducted according to European Directive 2010/63 (Recommendation Number 9). The study was approved under statement A15-035 from the “Comité d’Ethique en Expérimentation Animale du CEA” and was registered and authorized under National Number APAFIS #27263-2020091814549436 v2 by the French Ministry of Education and Research. Animals were observed daily after infection to check for any clinical signs or abnormal behavior, such as stress, abnormal interactions with staff, changes in feeding patterns, diarrhea, dyspnea, and cough. Experimental procedures (animal handling, bacterial exposure, and sampling) were performed under anesthesia with a combination of ketamine (5 mg/kg) and medetomidine (0.05 mg/kg). During anesthesia, pulse oximetry was performed and respiratory rate, heart rate, and weight were measured to calculate the clinical score and evaluate human endpoints. Animals were sacrified after with a bolus of sodium pentobarbital (180 mg/kg iv) after anesthesia and their tissues were collected for analysis.

### Animal infection

Animals were premedicated with atropine (0.04 mg/kg) 20 minutes before anesthesia with ketamine (5 mg/kg) and medetomidine (0.05 mg/kg). Macaques were intubated, and anesthesia was maintained with 1% isoflurane and oxygen, with pulse oximetry and cardiovascular parameters monitored during the procedure. A 2.8 mm flexible bronchoscope (Karl Storz) was inserted into the endotracheal tube and guided to the right caudal lung lobe by a trained veterinary surgeon. The macaque was infected by depositing 2 ml of a bacterial suspension containing 25 colony forming units (CFU) of *Mtb* Erdman via the bronchoscope operating channel, as previously described(23–25). Anne Lenaerts from Colorado State University kindly provided Roland Brosch at the Institut Pasteur in Paris, France with the *Mtb* Erdman strain. The inoculum was tittered after each exposure to ensure the correct bacterial load, and a CT scan performed four weeks after exposure confirmed the presence of lung granuloma in the targeted area.

### Serial positron emission tomography/computed tomography (PET/CT) imaging and image analysis

Lung lesions were monitored by positron emission tomography-computed tomography (PET/CT) imaging with the Digital Photon Counting (DPC) PET-CT system (Vereos-Ingenuity, Philips) implemented in an animal biosafety level 3 facility(45). An initial imaging session was performed before infection. After bacterial exposure, images were acquired every two weeks until the 12^th^ week and monthly thereafter, until the end of the study. Macaques were anesthetized with ketamine (5 mg/kg) and medetomidine (0.05 mg/kg), intubated, and placed on a warming blanket (Bear Hugger, 3M) on the machine bed. During the experiment, animals were maintained under anesthesia with 0.5-1.5% isofluorane in oxygen, and their vital parameters (cardiac rate, oxygen saturation, and temperature) were monitored. Lung CT images were acquired with a 64 x 0.6 mm detector collimation, a tube voltage of 120 kV, and an intensity of approximately 150 mA. Images were reconstructed with a slice thickness of 1.25 mm and an interval of 0.75 mm. A whole-body PET scan (5 steps, 3 min/step) was performed approximately 40 minutes after the intravenous injection of fluorodeoxyglucose ([^18^F]FDG) (3.13 ± 0.19 MBq/kg). PET images were reconstructed on a 256 x 256 matrix, with OSEM (3 iterations, 15 subsets). PET and CT images were analyzed with 3D slicer (open-source tool) software to determine the volume of FDG-positive lung lesions and the maximum radioactive signal for tracheobronchial lymph nodes.

### Determination of bacteria load in tissues and bronchoalveolar lavages

Bronchoalveolar lavage (BAL) was performed with a 2.8 mm flexible bronchoscope (Karl Storz) inserted into the endotracheal tube and guided to the right caudal lung lobe by a trained veterinary surgeon. The lung lobe was washed three times with 15 ml of saline per wash, via the operating channel. At necropsy, the tissues harvested (lung lobes, lymph nodes, spleen and liver) were mechanically digested with the gentleMACS dissociator system (Miltenyi Biotec Inc.) to obtain homogenates. Serial dilutions of BAL fluid and tissue homogenates were plated on 7H11 agar medium supplemented with 10% OADC (oleic acid-albumin-dextrose-catalase (BD)) and the PANTA (polymyxin B, amphotericin B, nalidixic acid, trimethoprim and azlocillin, (BD)) antibiotic mixture and incubated at 37°C. Colony forming units (CFUs) were counted after three weeks of incubation to determine the numbers of viable bacilli in each of the samples.

### Interferon gamma release assay: ELISpot

The IFN-gamma response to *Mycobacterium tuberculosis* was determined weekly throughout the study, with the Monkey IFN-γ ELISpot PRO kit (Mabtech Monkey IFN-γ ELISpot pro), in accordance with the manufacturer’s instructions and as previous described(46). Briefly, peripheral blood mononuclear cells (PBMC) were isolated from heparin-anticoagulated blood samples from macaques by the Ficoll® Paque density gradient centrifugation method. The suspensions of PBMC obtained were dispensed in the wells of a plate in duplicate. The cells were stimulated with ESAT-6, CFP-10 or PMA/ionomycin (Sigma Aldrich) and incubated for 18 h at 37°C under an atmosphere containing 5% CO_2_. Biotinylated anti-IFN-γ antibody was added, followed by ready-to-use BCIP/NBT-plus substrate solution that had been passed through a filter with 0.45 µm pores, and the plates were incubated. The ELISPOT plates were read with an Automated ELISpot Reader ELR08IFL (Autoimmun Diagnostika GmbH, Strassberg, Germany) and the number of spots per well, spot size and signal intensity per spot were analyzed.

### RNA extraction and sequencing

We collected 0.5 ml peripheral whole blood from each of six macaques. These samples were stored in Tempus medium at -80 °C. RNA was extracted with the Tempus Spin RNA Isolation Reagent Kit (Applied Biosystems). Sample concentration was determined with the Qubit™ RNA HS Assay Kit (Thermo Fisher Scientific). RNA quality was assessed with High-Sensitivity RNA ScreenTape and reagents (Agilent Technologies) on a TapeStation 4150 instrument (Agilent Technologies). In total, 100 ng of RNA per sample was denatured at 65°C and reverse-transcribed by a strand-switching technique using the Maxima H Minus Reverse Transcriptase (Thermo Fisher Scientific) to synthesize a double-stranded cDNA. PCR, barcoding, and adapter attachment were performed with the PCR-cDNA Sequencing Kit (SQK-PCB109, Oxford Nanopore Technologies). Samples were quantified with the QuBit 1x dsDNA HS kit (Thermo Fisher Scientific) and pooled in equimolar ratios before loading onto R9.4.1 Flow cells with the GridION instrument (MinKNOW version 22.08.9 or 22.10.5).

### RNA-seq data processing and analysis

Sequence reads were converted into FASTQ files. The FASTQ sequences associated with this project have been deposited in the SRA under BioProject ID PRJNA1236326 (http://www.ncbi.nlm.nih.gov/bioproject/1236326). Reads with a quality score below 9 were discarded. The remaining reads were aligned with both the *Macaca fascicularis* 5.0 and *Homo sapiens* GRCh38.p14 reference transcriptomes (GenBank assembly accession number GCA_000364345.1 and GCA_000001405), with minimap2 version 2.24 (H. Li, 2018). The mean of the median alignment ratio of reads on references in all samples was 0.86 for macaque and 0.87 for humans. The mean of median percent of read identities after alignment against the reference for all samples was 95% for macaque and 91% for human alignments. In addition, the proportion of unaligned reads with the macaque reference sequence reached 17%, whereas the proportion of unaligned reads with the human reference sequence was only 11% of total reads. Given that the *Macaca fascicularis* 5.0 reference genome has been removed from the National Center for Biotechnology Information (NCBI) databases for genome annotation processing and that read alignments with the human transcriptome were satisfactory, we decided to pursue quantification exclusively with the human alignment. For transcript quantification, the alignments obtained were processed with Salmon version 1.8.0 (Patro et al., 2017). For the exploration of differentially expressed genes, replicate count data were analyzed in R 4.3.0 with the DESeq2 package version 1.40.1. Both upregulated and downregulated genes were used for Gene Ontology (GO) Gene Set Enrichment Analysis (GSEA) and an analysis of overrepresentation with the Reactome database available from Reactome.org. GSEA was performed with clusterProfiler (version 4.9.0) to identify pathways from three GO collections that were over- and underrepresented: biological processes (BP), cellular components (CC) and molecular functions (MF). Normalized enrichment scores (NES > |1.5|) and false discovery rates (FDR < 0.05) were used to quantify the extent of enrichment and its statistical significance, respectively, and the enrichplot R package (version 1.20.0) was used for visualization. Unique genes for GO pathways of interest (GO: 0042742, defense response to a bacterium, and GO: 1904724, tertiary granule lumen) were selected. Heatmaps were generated with ComplexHeatmap package version 2.16.0.

### Cell isolation from whole blood, bone marrow (BM), bronchoalveolar lavage (BAL) and tissues for mass cytometry

Whole blood, bone marrow (BM), and BAL samples were collected before and after infection, as indicated in supplementary figure 1. Blood was collected in lithium-heparin tubes (Vacutainer BD, USA) by venipuncture. We harvested 1 to 2 ml BM were from the long bones with a Malarme trocar and deposited it in a Falcon 15 tube containing citrate dextrose solution. Cells were washed with 1X Dulbecco’s phosphate-buffered saline (DPBS) (Thermo Fisher Scientific), treated with 1X LGR to obtain erythrocyte-free leukocytes, and washed once with DPBS and once with RPMI 1640 medium (Thermo Fisher Scientific). At specific time points, BAL fluid was collected from the caudal right lobe by bronchoscope-guided lavage with three washes of 15 ml 0.9% NaCl. The blood, BM, and BAL were subjected to centrifugation (800g) and the pellet was resuspended in RPMI medium for staining for mass cytometry. On necropsy, lung granulomas, spleen, and tracheobronchial lymph nodes (LN) were collected in RPMI medium supplemented with 10 mM N-2-hydroxyethylpiperazine-N’-2-ethanesulfonic acid (HEPES) buffer (Sigma-Aldrich). Single cells were obtained from the lung granulomas by a combination of enzymatic and mechanical digestion. The lung granulomas were digested with collagenase D and DNAse I (Sigma-Aldrich) at 37°C for 30 minutes with the gentleMACS dissociator system (Miltenyi Biotec Inc). Splenocytes were isolated by physical digestion, based on gentle crushing on a cell strainer with a pestle. The cell suspensions obtained from digested tissues were gently filtered through a 70 µm-mesh nylon cell strainer (Falcon, BD Biosciences). Erythrocytes, if present, were removed from the filtrate preparations with red blood cell lysis buffer (1X LGR). The resulting cell suspensions were washed twice with RPMI medium, subjected to centrifugation at 800g, and the pellet was resuspended in RPMI medium for staining for mass cytometry.

### Mass cytometry immunophenotyping

Immunophenotypic surface and intracellular staining for mass cytometry was performed on whole blood, BM, BAL fluid, and tissue cell suspensions. We stained blood, BM, BAL, and tissue cell suspensions (5 × 10^6^ cells for each) with rhodium intercalator (viability dye) for 15 minutes at 37°C. The cells were washed three times with Maxpar staining buffer (Fluidigm, San Francisco, CA, USA), and stained with the heavy metal-labeled extracellular antibodies indicated in Supplementary table 1 (blue) for 20 minutes at room temperature. Before adding the extracellular antibodies, we saturated the tissue cells by incubation with 10% macaque serum. The stained cells were washed, fixed with 1.6% paraformaldehyde (PFA)/DPBS, and permeabilized with perm buffer (Invitrogen, Waltham, MA, USA). The cells were then stained with the heavy metal-labeled intracellular antibodies indicated in STAR protocol (red) for 30 minutes at room temperature. Three washes were performed with 1X DPBS. The cells were resuspended in 1.6% PFA/DPBS supplemented with 0.1 µM iridium nucleic acid intercalator and stored overnight at 4°C. The cells were washed three times with Maxpar water (Fluidigm, San Francisco, CA, USA), resuspended in an appropriate volume of Maxpar water, and passed through a 35 μm-mesh nylon cell strainer (BD Biosciences). EQTM Four-Element calibration beads (Fluidigm, San Francisco, CA, USA) were added to the cell suspension according to the manufacturer’s instructions, and the cells were acquired on a Helios machine (Fluidigm, San Francisco, CA, USA). All the samples were acquired on the same day to minimize instrument-related variation.

### Mass cytometry data processing and analysis

Mass cytometry data were normalized and concatenated (when necessary) with Fluidigm CyToF software (Fluidigm, San Francisco, CA, USA). Data abnormalities due to issues with the instruments, samples and acquisitions were removed with OMIQ® flowCUT algorithms. The samples were manually gated on the OMIQ platform to remove debris, normalization beads, dead cells, and doublets for the identification of CD45^+^ cells for subsequent analysis. We exported the data for CD45^+^ cells to R statistical software and transformed them by Arcsinh transformation (cofactor = 5). The data were clustered with flowSOM, a self-organizing map-based algorithm, with the som function of the kohonen package and a som grid size of 20 x 20 (blood), 28 x 28 (BAL, BM, lung and lymph node), or 25 x 25 (spleen). As additional parameters, we used rlen = 100, mode = pbatch. The metaclustering step was performed with the hclust function and the Ward criterion. The data underwent a reduction of dimensions and were visualized by uniform manifold approximation projection (UMAP) with the umap function from the uwot package and the following parameters: init = laplacian, learning_rate = 0.5, n_epochs = 500, n_neighbors = 25, and min_dist = 0.01. Automated optimized parameters for T-distributed stochastic neighbor embedding (opt-SNE) were computed for blood neutrophils with the tsne function of the Rtsne package and the following markers: CD64, HLADR, CD66abce, CD32, CD69, PD1, CD163, CD10, CD11b, CD45RA, CD95, CD14, CD62L, CD101, CCR5, CD28, NKg2a, CD16, CD11c, FceRIa CD45. The additional parameters used were initial_dims = 10, perplexity = 30, theta = 0.5, and max_iter = 5000.

### Isolation of neutrophils for NET and ROS determinations

We collected 6 ml peripheral blood in lithium-heparin tubes (Vacutainer BD, USA) and removed the PBMC by Ficoll-Paque density gradient separation. The pellets containing the granulocytes were treated with red blood cell lysis buffer (LGR 1X) to eliminate erythrocytes. The erythrocyte-free cell preparation was then washed and resuspended in RPMI medium.

### Determination of neutrophil extracellular trap (NET) levels

We suspended 1 × 10^6^ cells in 200 μl NET assay medium (RPMI medium, 10 mM HEPES and 1% macaque serum) and left them unstimulated or stimulated them with 100 nM phorbol myristate acetate (PMA) (Sigma-Aldrich) or 200 μg *Staphylococcus aureus* peptidoglycan (Sigma-Aldrich) at 37°C under an atmosphere containing 5% CO_2_ for 4 hours. We added 1 μg DNAse I (Sigma-Aldrich) and incubated for 15 minutes at room temperature to achieve the partial digestion of NETs. The reaction was stopped by adding 5 mM ethylenediaminetetraacetic acid (EDTA). The cells were vigorously mixed with a pipette, centrifuged at 300 x *g* for 5 minutes at room temperature. The supernatant was carefully removed by aspiration and stored at -80°C. We coated 96-well ELISA plates were coated with anti-MPO antibody at a final concentration of 20 μg/mL by incubation overnight (18 hours) at 4°C.

The plates were washed with DPBS 1X-0.05% Tween-20 and non-specific binding was blocked by incubation with blocking buffer (2% BSA in 1xDPBS) for 2 hours at room temperature. The plates were washed three times and a 1/16 serial dilution of the sample was added to the well and incubated for 1 hour at room temperature. The plates were washed three times, the horse-radish peroxidase (HRP) conjugated anti-DNA secondary antibody (Sigma-Aldrich) was added and the plates were incubated for one hour at room temperature. The wells were washed again, the substrate 2,2’-azino-bis-3-ethylbenzothiazoline-6-sulfonic acid (ABTS) was added, and the plates were incubated for 40 minutes in the dark at room temperature. The reaction was then stopped by adding the stop solution provided in the cell death kit, and absorbance was measured with a microplate reader (TECAN Spark). The NETosis ratio was calculated as follows: 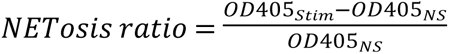

### Determination of reactive oxygen species (ROS)

We suspended 5 × 10^5^ cells in 200 µl ROS assay medium (RPMI medium, 10 mM HEPES and 1% macaque serum). N-acetylcysteine (NAC, Thermo Fisher Scientific) was added for the negative control, and the plates were incubated for 1 h at 37°C. The cells were then either left unstimulated or were stimulated with Tert-butyl hydroperoxide (t-BHP) (Thermo Fisher Scientific) for the positive control, or with 100 ng/ml PMA (Sigma-Aldrich), 10 nm fMLP (Sigma-Aldrich), 10^7^ heat-killed Mtb or 10^6^ Mtb lysate for 30 minutes at 37°C. CellROX green dye (Thermo Fisher Scientific) was added to the cells, which were incubated for a further 15 minutes at 37°C. The antibody mixture (antibodies against CD45, CD66abce, CD10, CD101 and CD62L) for cell phenotyping was added and the cells were incubated for a further 15 minutes at room temperature. The cells were washed with 1xDPBS, centrifuged, and the pellet was resuspended in 4% PFA in 1X DPBS and subjected to acquisition on a ZE5 flow cytometer (Bio-Rad) equipped with five lasers (355 nm ultraviolet, 405 nm violet, 488 nm blue, 561 nm yellow and 640 nm red lasers).

### Neutrophil-T cell coculture assay

PBMC and granulocytes were isolated from whole blood by Ficoll-Paque density gradient centrifugation. Erythrocytes were eliminated from the two cell populations by incubation with red blood cell lysis solution (LGR 1X). We enriched the granulocyte fraction in neutrophils by performing magnetic depletion to remove non-neutrophil cells. Briefly, cells were stained with anti-CD125 antibodies conjugated with PE (Miltenyi Biotec) and washed with AutoMACs Buffer (Miltenyi Biotec). Anti-PE and anti-CD2 microbeads (Miltenyi Biotec) were added to the granulocyte fraction to target eosinophils and the remaining CD2^+^ cells, respectively, and magnetic depletion was performed on LD columns (Miltenyi Biotec). This technique resulted in a neutrophil purity >95%. Isolated PBMC were stimulated (TCR stimulation with Cytostim NHP, Miltenyi Biotec) and cultured in the presence or absence (0:1) of autologous neutrophils at ratios of 1:1, 3:1 and 5:1 neutrophils to PBMC, together or separately, in 0.4 µm-pore 96-well Transwell plates (Corning) in the presence of costimulatory antibodies (directed against CD28/CD49d, BD Bioscience) and an anti-CD107a antibody (Miltenyi Biotec). The cells were incubated for 1 h at 37°C under an atmosphere containing 5% CO_2_. Brefeldin A (10 µg/ml, Sigma-Aldrich) and Monensin (Biolegend) were added to the wells, and the cells were incubated overnight at 37°C. They were then washed with PBS and viability staining was performed with Live/Dead (Thermo Fisher Scientific) stain for 20 min at 4°C. The cells were washed and permeabilized/fixed with Cytofix/Cytoperm reagent (BD Bioscience). They were then stained with the antibodies indicated in Supplementary table 3 for 30 minutes at 4°C. The stained cells were washed, suspended in fixation buffer (Biolegend) and acquired on a ZE5 flow cytometer (Bio-Rad) equipped with five lasers. The percent increase was calculated as follows:

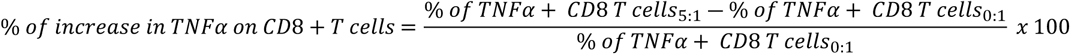

### Necropsy

^18^F-FDG PET/CT imaging was performed before necropsy, to assess disease progression and identify identify individual granulomas. This PET/CT examination identified lung lesions (granulomas), and lesions that not otherwise visible, and the lymph nodes were harvested. The location, size and estimated time of granuloma formation were determined by prenecropsy scans and during necropsy. Sample tissues from the spleen, liver and other apparent extrapulmonary lesions were harvested, as previously described. Harvested lesions were divided into samples for histology, immunohistochemistry, mass cytometry and determinations of bacterial load.

### Histology and immunohistochemistry (IHC)

The tissues harvested during necropsy were fixed by immersion in 10% formalin and embedded in paraffin. Thin sections were cut and stained with hematoxylin and eosin (H&E) and calprotectin-DAB stain for neutrophil identification by standard procedures. We imaged slides with an Axioscan 7 Scanner (Zeiss). Calprotectin-positive cells were segmented and counted with QuPath in different parts of the tuberculous granuloma and lung parenchyma.

### Statistical analysis

Statistical analysis was performed with GraphPad Prism version 10.0, R packages and SRPlot. Comparisons between two groups were performed with the non-parametric Wilcoxon and Mann-Whitney *U* test. Comparisons between two groups and different types of points were performed with Kruskal-Wallis tests with Tukey’s correction. All the details of the statistical analysis, including the type of test used can be found in the legends to the figures. Data are presented as the mean ± SEM (standard error of the mean). \**p* < 0.05, \*\**p* < 0.01, \*\*\**p* < 0.001, ns = not significant. For transcriptomic analysis, adjusted *p*-values were used and false discovery rates were considered.

## Resource availability

Requests for further information and resources should be directed to and will be satisfied by the lead contact, Julien Lemaitre

## Availability of materials

This study generated no new unique reagents.

## Data and code availability

The whole-blood transcriptomic FASTQ sequences associated with this project have been deposited in the SRA under BioProject ID PRJNA1236326 and are publicly available as of the date of publication, from http://www.ncbi.nlm.nih.gov/bioproject/1236326.

## Author contributions

Conceptualization, S.B.D., C.J, B.D, M.H, R.L.G. and J.L.; Methodology, S.B.D., C.J, V.M., N.N., C.M., M.L., G.S., S.L., B.D., W.Z., R.B., F.R., Q.P., B.J., M.H., T.N. and J.L.; Software, P.M. and N.N.; Validation, C.J., N.N., A.S.G., N.B., F.R., T.N., R.L.G. and J.L.; Formal analysis, S.B.D, C.J., P.M., N.N., S.D., G.S., S.L., E.J., N.B., Q.P., B.J., M.H., T.N. and J.L.; Investigation, S.B.D, C.J., V.M., N.N., C.M., G.S., J.M., S.L., C.L., B.D., E.J., W.Z., B.J., M.H. and J.L.; Resources, W.Z., R.B., V.C., A.S.G., N.B., F.R., O.L., T.N. and R.L.G.; Data curation, C.J., P.M., N.N., N.B. and J.L.; Writing – original draft, S.B.D., C.J., T.N., R.L.G. and J.L.; Writing – Review & Editing, S.B.D., C.J., N.N., B.D., W.Z., R.B., Q.P., T.N., R.L.G. and J.L.; Visualization, S.B.D., C.J., P.M., N.N., C.M., G.S., Q.P., T.N. and J.L.; Supervision, C.J., A.S.G., N.B., R.R., Q.P., M.H., O.L., T.N., R.L.G. and J.L., Project administration, C.J., V.C., R.L.G. and J.L., Funding acquisition, C.J. and R.L.G.

## Acknowledgments

We thank all members of the Animal Science and Welfare, FlowCyTech, Imaging of Infection and Immunity, Immunology and Infectiology core facilities of the IDMIT infrastructure for their excellent expertise and outstanding contributions: Quentin Sconosciuti, Maxime Potier, Eleana Navarre, Jean-Marie Robert, Emma Burban, Sebastien Langlois, Julie Morin, Kyllian Lheureux, Matthieu Van Tilbeurgh, Wesley Gros, Ernesto Marcos, Laetitia Bossevot, Maxence Galpin, Loïc Pintore, Sylvie Legendre, Yann Gorin, Alicia Pouget, and Sebastien Jacquin. We thank our collaborators from Oncodesign services: Julien Dinh, Eloise Joffroy, Alexandre Baillet and Elodie Guyon. We also thank Fadel Sayes, and Alexandre Pawlik from the Institut Pasteur Paris for their methodological advice regarding Mtb strain handling. We also thank Luc de Chaisemartin for sharing his expertise on neutrophils.

## CORRESPONDING AUTHORS’ CONTACT INFORMATION

Julien Lemaitre, Université Paris-Saclay, INSERM, CEA

U1184 IMVA-HB/IDMIT, Fontenay-aux-Roses, France

Roger Le Grand, Université Paris-Saclay, INSERM, CEA

U1184 IMVA-HB/IDMIT, Fontenay-aux-Roses, France

